# GWAS reveals the genetic complexity of fructan accumulation patterns in barley grain

**DOI:** 10.1101/2020.06.29.177881

**Authors:** Andrea Matros, Kelly Houston, Matthew R. Tucker, Miriam Schreiber, Bettina Berger, Matthew K. Aubert, Laura G. Wilkinson, Katja Witzel, Robbie Waugh, Udo Seiffert, Rachel A. Burton

**Author notes:** **Corresponding author:** Andrea Matros, ARC Centre of Excellence in Plant Energy Biology, School of Agriculture, Food and Wine, University of Adelaide, Adelaide, South Australia, Australia. **E-Mail addresses:**.

## Abstract

We profiled the grain oligosaccharide content of 154 two-row spring barley genotypes and quantified 27 compounds, mainly fructans, that exhibited differential abundance. Clustering revealed two major profile groups where the ‘high’ set contained greater amounts of sugar monomers, sucrose and overall fructans, but lower fructosylraffinose. GWAS identified a significant association for the variability of two fructan types; neoseries-DP7 and inulin-DP9 which showed increased strength when a compound-ratio GWAS was applied. Gene models within this region included five fructan biosynthesis genes, of which three (*fructan:fructan 1-fructosyltransferase*, s*ucrose:sucrose 1-fructosyltransferase*, and s*ucrose:fructan 6-fructosyltransferase)* have already been described. The remaining two, *6(G)-fructosyltransferase* and *vacuolar invertase1* have not previously been linked to fructan biosynthesis in barley and showed expression patterns distinct from those of the other three genes, including exclusive expression of *6(G)-fructosyltransferase* in outer grain tissues at the storage phase. From exome capture data several SNPs related to inulin- and neoseries-type fructan variability were identified in *fructan:fructan 1-fructosyltransferase* and *6(G)-fructosyltransferase* genes Co-expression analyses uncovered potential regulators of fructan biosynthesis including transcription factors. Our results provide evidence for the distinct biosynthesis of neoseries-type fructans during barley grain maturation plus new gene candidates likely involved in the differential biosynthesis of the various fructan types.

**Highlight:** Grain fructan profiles in barley are more complex than previously expected and variations in a diversity panel relate to a genomic region where fructan biosynthesis genes cluster.

## Introduction

Starch, fructans and (1,3; 1,4)-β-glucans represent the major plant reserve carbohydrates (Vijn and Smeekens, 1999; Burton and Fincher, 2009). Among them, fructan biosynthesis has evolved polyphyletically in about 15% of higher plants, including species of the orders Asterales, Buxales, Asparagales and Poales (Hendry and Wallace, 1993; Cairns *et al.*, 2000; Van den Ende, 2013). In cereals, fructans accumulate in all plant organs (Pollock and Cairns, 1991).

Fructans consist of repeating fructose residues linked to a sucrose unit. The classification relates to the position of the sucrose, the linkage-type between the fructose residues (i.e. ß(2,1), inulin; ß(2,6), levan; or containing both ß(2,1) and ß(2,6)-d-fructosyl units referred to as graminan-type) and the chain lengths (Cochrane, 2000; Matros *et al.*, 2019). Fructans can form oligomers with a degree of polymerization (DP) of 3-9 or polymers with a DP ≥10. Here, fructans is used to indicate either fructooligosaccharides (FOS) or fructan polymers. Fructans are typically discussed in the literature without differentiation of the DP, but since they have become more important in a dietary context (Dwivedi *et al.*, 2014; Verspreet *et al.*, 2015b; Liu *et al.*, 2017) more attention has recently been paid to the role of fructans according to their DP level.

All types of fructans are known to occur in the Poaceae (Carpita *et al.*, 1991; Pollock and Cairns, 1991; Bonnett *et al.*, 1997). However, *Triticum*, *Secale* and *Hordeum* are believed to mainly contain branched-type fructans (graminan-type) whereas the Poeae tribe mostly comprises levan-type fructans (Bonnett *et al.*, 1997; Huynh *et al.*, 2008a). Recently, the presence of graminan- and neoseries-type fructans was reported in wheat (Verspreet *et al.*, 2015c). Neoseries-type fructans, in contrast to other fructan-types, are characterised by an internal glucose unit (Matros *et al.*, 2019). Additional structural variations are likely to occur between different plant organs.

New developments in fructan analysis based on mass spectrometry (MS) detection revealed the fine structure of cereal grain fructans with DP3-5 (Verspreet *et al.*, 2017). Variations in fructan composition pattern and abundance were observed in oat, barley, rye, spelt and wheat flour, suggesting a putative link between accumulation of certain fructan types and cereal phylogeny (Verspreet, *et al.*, 2017).

Reports of the beneficial health effects of fructans (Verspreet *et al.*, 2015b; Liu *et al.*, 2017; Anrade *et al.*, 2019) have prompted screens for variation in their natural abundance and composition and biotechnological approaches to increase FOS content in classical non-fructan cereals, such as maize (Dwivedi *et al.*, 2014). However, most studies on grain fructan content still focus on wheat (Huynh *et al.*, 2008a and b; Veenstra *et al.*, 2017; Veenstra *et al.*, 2019). Investigation of two doubled haploid (DH) populations (Berkut x Krichauff and Sokoll x Krichauff) revealed several quantitative trait loci (QTL) for high fructan content in wheat grain (Huynh *et al.*, 2008b). Winter wheat grain fructan content was found to be significantly influenced by either the genotype or the environment as well as by genotype × environment interactions (Veenstra *et al.*, 2019). Fructan content in developing barley grain was compared between seven genotypes, demonstrating peak accumulation between 6 and 17 days after pollination (DAP) (De Arcangelis *et al.*, 2019) as previously reported (Peukert *et al.*, 2014). Notably, a comparative mapping approach involving wheat and barley revealed clusters of genes encoding fructan biosynthesis enzymes (Huynh *et al.*, 2012) on 7AS in wheat and 7HL in barley. These clusters included *sucrose:sucrose 1-fructosyltransferase* (*1-SST*), *fructan:fructan 1-fructosyltransferase* (*1-FFT*), *sucrose:fructan 6-fructosyltransferase* (*6-SFT*), and several vacuolar invertases. Similar gene structures and physical positions of these clusters of functionally related genes in both genomes indicate that they may have evolved in parallel and that the genes within a cluster may be linked functionally in controlling fructan accumulation.

Due to its increasing potential as a health-promoting functional cereal, there is considerable interest in identifying factors that influence barley grain quality (Meints *et al.*, 2016; Langridge and Waugh, 2019). Here we report on an analysis of natural variation in fructan content and composition across a diversity panel of two-row spring barley. We identified significant associations between fructan composition/content and fructan biosynthesis genes. We obtained support for the involvement of some of these in underpinning the observed variation from transcriptomic analysis. Additionally, potential regulators of fructan biosynthesis were assigned by co-expression analyses.

## Materials and methods

### Plant material

We used 154 two-row spring barley genotypes sourced from The James Hutton Institute, complemented by three Australian elite barley varieties and the wheat line Piccolo as checks (Table S1). The germplasm was selected for minimum population structure while maintaining as much genomic diversity as possible based on principle components analysis of a much larger set of genotypes (>800). Three plants per genotype (biological replicates) were grown in a randomised main-unit design in a glasshouse compartment in a mix of clay-loam and cocopeat (50:50 v/v) and day/night temperatures of 22°C/15°C between July and December 2014 in The Plant Accelerator, Adelaide, Australia. Mature grains were harvested and stored until oligosaccharide analysis. For each sample, five grains were ground together to a fine powder using a PowerLyzerTM24 Homogenizer (QIAGEN) and used for oligosaccharide analysis immediately.

### Oligosaccharide extraction and profiling

A ‘mixed sample’ was assembled composed of equal amounts from each individual sample (154 genotypes x three biological replicates) to capture systematic shifts during extraction and measurement. Soluble sugars were extracted following a method adapted from Verspreet *et al.* (2012) by incubation in 80% ethanol at 85°C for 30 min followed by Milli-Q water at 85°C for 30 min on a mixer (700 rpm) in a final dilution of 1:40 (w/v, mg/μl), and supernatants combined. Extracts were diluted with water to 1:1000 (w/v, mg/μl) and 25 μl per sample analysed by high pH anion exchange chromatography with pulsed amperometric detection (HPAEC–PAD) on a Dionex ICS-5000 system using a DionexCarboPAC™ PA-20 column (3 x 150 mm) with a guard column (3 x 50 mm) kept at 30°C and operated at a flow rate of 0.5 ml min^−1^. The eluents used were (A) 0.1 M sodium hydroxide and (B) 0.1 M sodium hydroxide with 1 M sodium acetate. The gradient used was: 0% (B) from 0-2 min, 20% (B) from 2-35 min, 100% (B) from 35-36.5 min, 0% (B) from 37.5-38.5 min. Detector temperature was maintained at 20°C, data collection was at 2 Hz and the Gold Standard PAD waveform (std. quad. potential) was used.

Data acquisition, processing, and peak integration were performed using the Chromeleon™ version 7.1.3.2425 software (Thermo Scientific). Compounds were annotated based on available analytical standards. Glucose, fructose, sucrose, raffinose, 1-kestose, maltose, maltodextrin, nystose and mixtures of inulin from chicory (DP2-60) and levan from *Erwinia herbicola* were purchased from Sigma-Aldrich, while 1,1,1-kestopentaose was obtained from Megazyme. Additional inulin and neoseries-type fructans were isolated from onions and barley grain and analysed by mass spectrometry (MS). Fructan-related chromatographic peaks were identified based on fructanase digestion and mild acid hydrolysis (Supplementary Methods). A total of 27 peaks were annotated (Table S2).

### Metabolic data analyses

Peak area entry means and variances with respective standard deviations were calculated in Excel 2007 (Microsoft) from the ‘mixed sample’-normalised integrated peak area values of the individual biological replicates for each two-row spring barley line and the check lines (Table S3). Data from at least three biological replicates were available for 143 lines. For ten lines (Agenda, Alliot, Appaloosa, Cellar, Drought, Goldie, Scarlett, Tankard, Tartan, Turnberry), data from two biological replicates were available. The mean values for the two lines with just one entry (Calgary and Saana) were replaced by the only available data. Bonferroni outlier test was performed and pair-wise correlations between the abundances of the 27 metabolites were revealed by applying the average linkage clustering method, based on Pearson correlation coefficients implemented in the MVApp (Julkowska *et al.*, 2019; http://mvapp.kaust.edu.sa/MVApp/). The metabolite abundances were analysed with the software package MATLAB (The MathWorks, Inc.) with a log-logistic distribution applied. The Neural Gas (NG) algorithm, implemented in MATLAB was applied for cluster analysis following Kaspar-Schoenefeld *et al.* (2016) and Peukert *et al.* (2016). Analyses were performed for the biological replicates individually and the number of NG clusters was set to four.

### GWAS

GWAS was carried out by combing the phenotypic data for the 154 barley accessions with genotypic data generated using the Barley 50K iSelect genotyping platform (Bayer *et al.*, 2017). We focused on two-row spring barley accessions to reduce the confounding effects of population structure (Comadran *et al.*, 2012) that could have been introduced by including other row types and growth habits (Darrier *et al.*, 2019). Prior to analysis any single nucleotide polymorphism (SNP) with a minimum allele frequency (MAF) of < 0.05 was removed which left 24,925 polymorphic markers for our analysis. Marker-trait association analysis was carried out using R 2.15.3 (http://www.R-project.org) and performed with a compressed mixed linear model (Zhang *et al.*, 2010) implemented in the GAPIT R package (Lipka *et al.*, 2012). Linkage disequilibrium (LD) was calculated across the genome between pairs of markers using a sliding window of 500 markers and a threshold of R^2^ < 0.2 using Tassel v 5 (Bradbury *et al.*, 2007) to identify local blocks of LD, facilitating a more precise delimitation of quantitative trait loci (QTL) regions. We anchored regions of the genome containing markers that had passed the Benjamini-Hochberg threshold (p < 0.05) as implemented in GAPIT to the barley physical map (Mascher *et al.*, 2017) using marker positions provided in Bayer *et al.* (2017) and then expanded this region using local LD derived from genome wide LD analysis as described above. Putative QTL represented by less than 5 SNPs with −log10(p) values < 3 were not considered to be robust given the marker density and extensive LD present in the barley genome (Mascher *et al.*, 2017). The SNP with the highest LOD score was used to represent a significant QTL. We investigated significantly associated regions using BARLEX (https://apex.ipk-gatersleben.de/apex/f?p=284:39) to identify putative candidate genes. Gene annotations refer to entries in the UniProt database (https://www.uniprot.org/uniprot/, June 2019). Unknown genes were searched against the non-redundant entries for plants (June 2019) in the NCBI database using the BLASTX 2.9.0+ software (https://blast.ncbi.nlm.nih.gov/Blast.cgi).

For compounds showing an association that passes the false discovery rate (FDR) calculated in GAPIT, ratios between these and all other compounds quantified were generated. The ratios were log transformed and then used to carry out further GWAS. The ‘p-gain’, defined as the ratio of the lowest p-value of the two individual metabolites and the p-value of the metabolite ratio (Petersen *et al.*, 2012) was then calculated. A critical value for the p-gain was derived using B/(2*α), where α is the level of significance (0.05) and B the number of tested metabolite pairs. As we tested fifty-two pairs of compounds our critical value threshold was 5.2 x 10^2^.

Publicly available exome capture datasets (Mascher *et al.*, 2017) were used to identify potential causal polymorphisms in candidate genes. We only considered non-synonymous SNPs with less than 10% missing data across the set of germplasm to be informative.

### Gene transcript expression analyses of various developmental stages and tissues

Transcript abundance of genes of interest were measured in whole germinated grain (mean data from genotypes Navigator and Admiral) and isolated Navigator grain tissues from 0 to 96 hours after imbibition (hai). Aleurone tissues were divided into approximately thirds, with the proximal aleurone closest to the embryo (Betts *et al.*, 2019).

Data for seedling tissues (germinated embryo, root, and shoot) were obtained from the Expression Atlas organ dataset (https://www.ebi.ac.uk/gxa/home).

Data from epidermal strips (4 weeks after sowing, W4), roots (W4), inflorescences, rachis (W5), inflorescences, lemma, inflorescences, lodicule, dissected inflorescences, palea (W6), inflorescence (10 mm), and internode, as well as for senescing leaf were obtained from BARLEX (https://apex.ipk-gatersleben.de/apex/f?p=284:10) (Colmsee *et al.*, 2015). A developing anther dataset was obtained from Barakate *et al.* (2020) covering four anther stages (premeiosis, leptotene/zygotene, metaphase I to tetrad, pachytene/diplotene) and two meiocyte stages (leptotene/zygotene, pachytene/diplotene). Raw expression data were mapped against the transcriptome of barley (https://webblast.ipk-gatersleben.de/barley_ibsc/downloads/; merging high-confidence and low-confidence transcripts as well as isoforms) using Salmon v14.0 (Patro *et al.*, 2007). RNA-sequencing data were obtained from developing pistils at Waddington (W) stages W8, W8.5, W9, W9.5 and W10 (Wilkinson *et al*., 2019) and are shown as a mean value for five genotypes including Golden Promise (1x replicate per stage; Aubert *et al*., 2018), Salka, Wren, Forum and Gant (2x replicates per stage). In addition, RNA-sequencing data from individual pistil tissues including the nucellus, integuments, ovary wall, embryo sac, egg apparatus and central cell, antipodal cells, and chalaza were analysed from the Sloop genotype.

Gene transcript expression data for whole developing grain (from 7 to 20 DAP) minus the embryo were generated by RNA-sequencing and are shown as a mean value from 6 genotypes including Sloop (1x replicate per timepoint; Aubert *et al*., 2018), Alabama, Pewter, Extract, Taphouse, and Hopper (1x replicate per timepoint), while isolated developing grain tissues of interest including the pericarp, aleurone, sub-aleurone, and starchy endosperm were generated from medial sections at 7 to 25 DAP for the genotype Sloop (1x replicate per timepoint).

### Correlation analyses of gene transcript expression

Correlations among transcript abundance of fructan metabolism genes with other gene models from the QTL interval detected in the GWAS were evaluated for each of the RNA-sequencing datasets listed above, individually. Pair-wise correlations between the gene transcript expression levels were revealed by applying the average linkage clustering method, based on Pearson correlation coefficients implemented in the MVApp (Julkowska *et al.*, 2019).

## Results

### Grain oligosaccharide profiling revealed the abundance of fructans

HPAEC-PAD chromatograms of non-structural soluble carbohydrates from mature barley grain allowed for separation of monosaccharides, disaccharides, and oligosaccharides with a DP <15 (Table 1, Figure S1). Among the latter, we identified two raffinose family oligosaccharides (RFO) and three maltose-type oligosaccharides. Most compounds were found to be fructans (Table S2), including levan-, inulin-, graminan, and NS-inulin-types. A high abundance of fructans with DP3 and DP4 was observed in mature grain extracts. Oligosaccharide profiles were obtained and evaluated from all 154 two-row spring barley lines and four checks (Table S1) and integrated peak areas extracted for 27 compounds (Table 1). The resulting data matrix was used for analyses of oligosaccharide distribution, abundance variation, metabolite correlations, and GWAS.

**Table 1:**
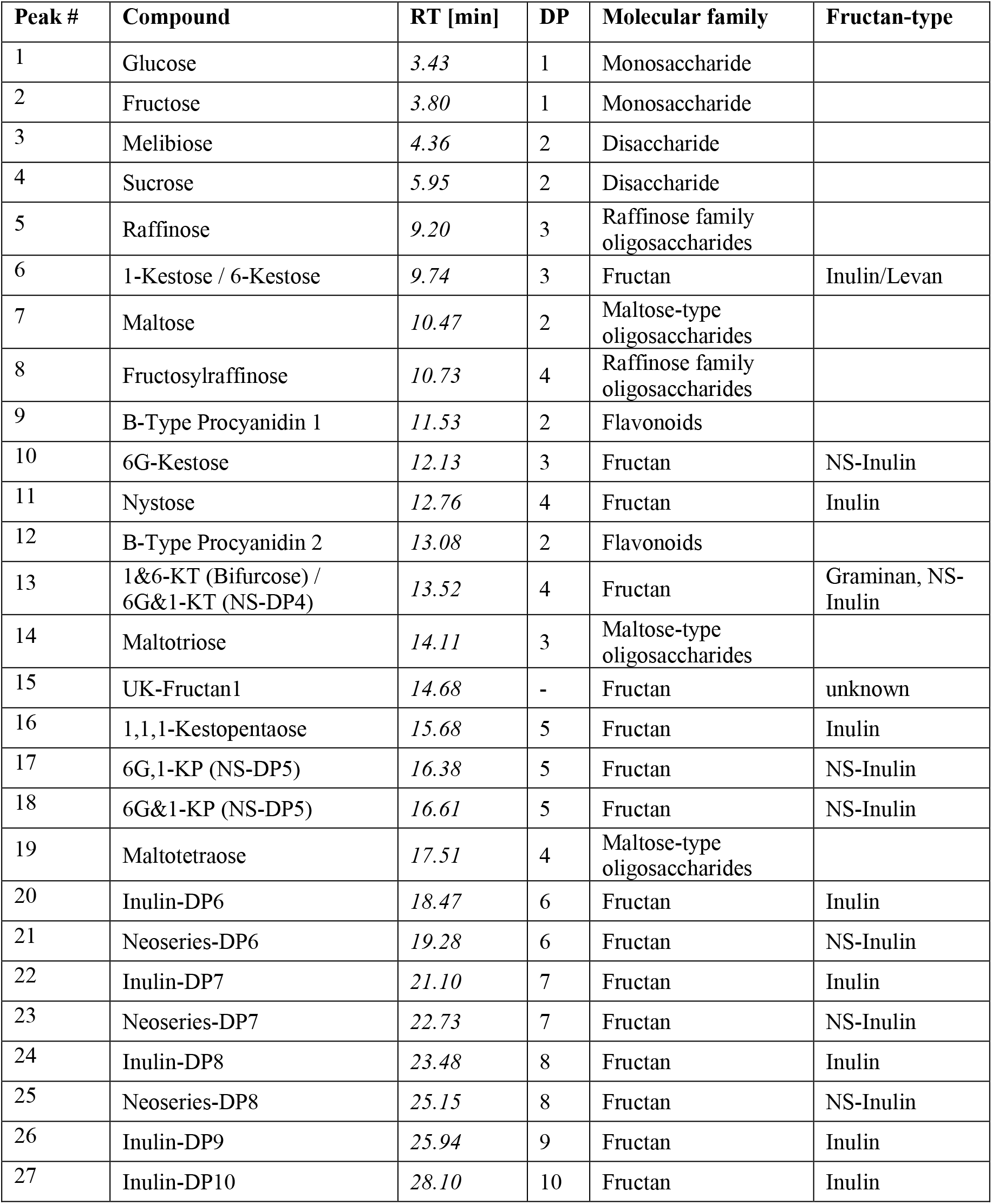
List of the 27 annotated metabolites. Peaks were annotated by comparison with analytical standards and isolated fractions from barley grain and onion bulb samples as well as based on fructanase digestion and mild acid hydrolysis. Further details of compound annotation are provided in Table S2. Abbreviations: DP, degree of polymerisation; KP, kestopentaose; KT, kestotetraose; NS, neoseries-type fructan; RT, retention time

### Large variations detected in oligosaccharide profiles

The abundance of most compounds followed a log-logistic distribution. Only the two most abundant compounds, sucrose and raffinose, followed a normal distribution (Figure S2). Oligosaccharide profiles were grouped separately for each of the biological replicates. Applying Neural Gas (NG) clustering to the data identified four statistically significant patterns of abundance (clusters). Each cluster can be interpreted as a prototypic abundance profile of the underlying metabolite values (Figure S3). They mainly differed in height of the normalised peak areas. Overlaps between cluster 1 and 4 and cluster 2 and 3 were detected. Samples in the latter clusters were characterised by significantly higher levels of sugar monomers and sucrose, lower fructosylraffinose and higher overall fructan values compared to clusters 1 and 4 samples (Figure S3). Accordingly, we rationalised the four clusters into two profile groups (Figure 1); cluster 1 and 4 forming profile group 1 (‘low’) and cluster 2 and 3 forming profile group 2 (‘high’). The largest peak in each sample was sucrose, whilst among the oligosaccharides, the highest values were detected for raffinose, the co-eluting fructans 1-kestose/6-kestose, nystose and the co-eluting 1&6-kestotetraose (KT, bifurcose)/6G&1-KT (NS-DP4). Generally, a higher abundance of fructans with DP3 and DP4 was observed for all accessions, and differentiation between individuals was mainly attributed to the overall abundance of all fructan types.

**Figure 1:**
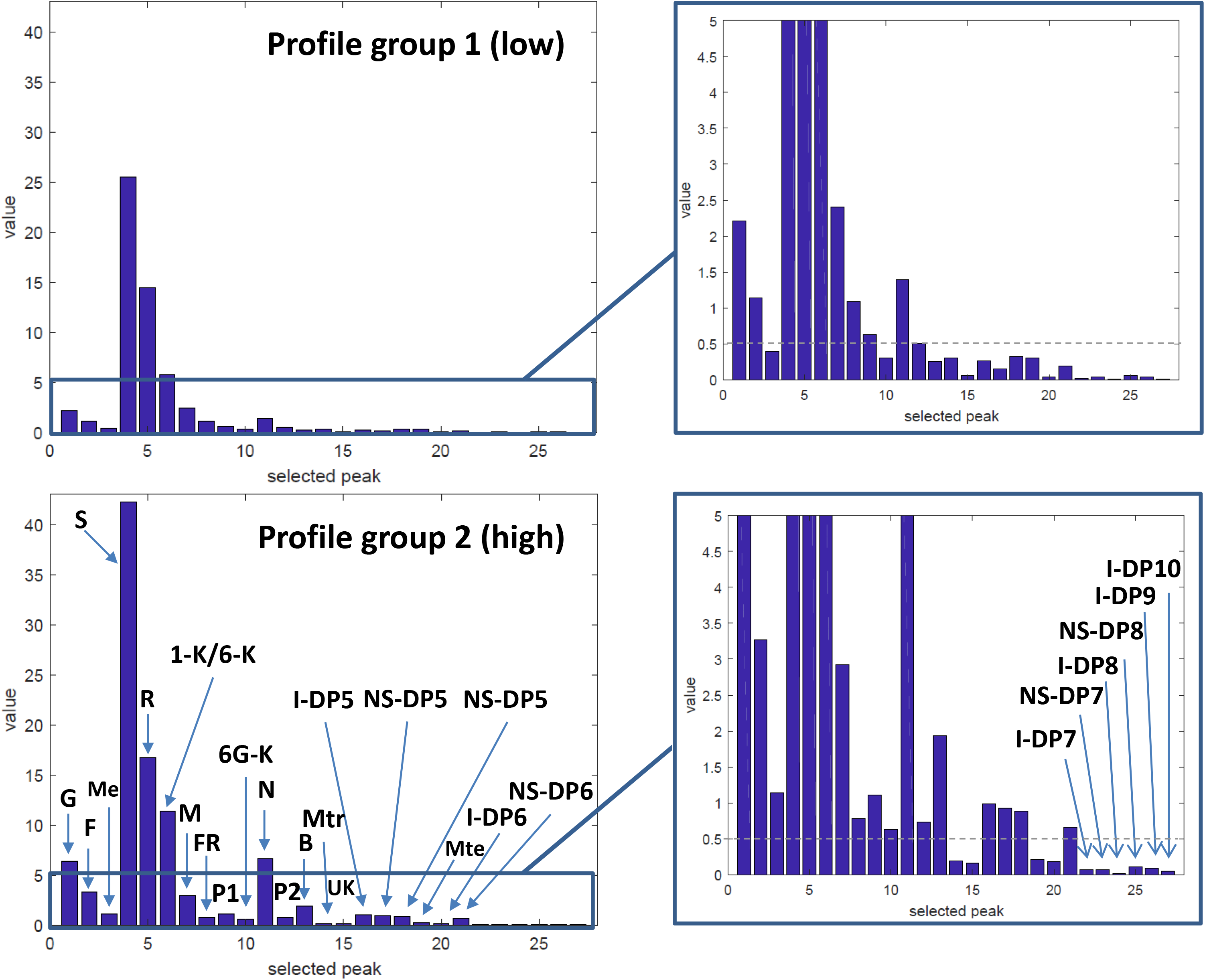
Oligosaccharide profile groups as obtained by Neural Gas clustering. Shown are the mean profiles of the two major profile groups. Value (y-axis) represents the peak area [nC*min] of the individual compounds (x-axis) listed in Table 1. Profile group 2 was characterised by higher levels of sugar monomers and sucrose, lower fructosylraffinose and higher overall fructan values. Abbreviations: G-glucose, F-fructose, Me-melibiose, S-sucrose, R-raffinose, 1-K/6-K-1-kestose/6-kestose, M-maltose, FR-fructosylraffinose, P1-procyanidin B1, 6G-K-6G-kestose, N-Nystose, P2-procyanidin B2, B-1&6-KT (Bifurcose)/6G#x0026;1-KT (NS-DP4), Mtr-maltotriose, UK-unknown fructan, I-DP5-inulin DP5, NS-DP5-neoseries-DP5 (probably 6G,1-KP and 6G&1-KP), Mte-maltotetraose, I-DP6-inulin DP6, NS-DP6-neoseries-DP6, I-DP7-inulin DP7, NS-DP7-neoseries-DP7, I-DP8-inulin DP8, NS-DP8-neoseries-DP8, I-DP9-inulin DP9, I-DP10-inulin DP10.

We then assigned individual barley accessions to profile groups according to abundance profiles in each individual replicate (Table S1). Accessions with only two biological replicates and mixed representation of their clusters in the two profile groups were assigned as ‘mixed’, as their oligosaccharide profile group was not distinct. In total 76, 77, and 5 accessions were assigned to the profile groups ‘low’, ‘high’, and ‘mixed’ respectively.

### Significant correlations are observed between metabolites

Of the 349 pair-wise correlations, 184 (52.72%) were highly significantly correlated (p < 0.001), 204 (58.45%) moderately significantly (p < 0.01) and 228 (65.33%) were just significant (p < 0.05) (Table S4). Several regions with highly correlated metabolites were identified in the results matrix, reflecting in many cases, biochemical relationships (Figure 2). The most significant positive correlations were observed between the various branched neoseries-type fructans as well as between the linear inulin-type fructans. Significant positive correlations were also detected between the monosaccharides and their related disaccharides as well as between the maltose-type oligosaccharides. The most significant negative correlations were detected between fructosylraffinose and fructans being highest with nystose (Figure 2).

**Figure 2:**
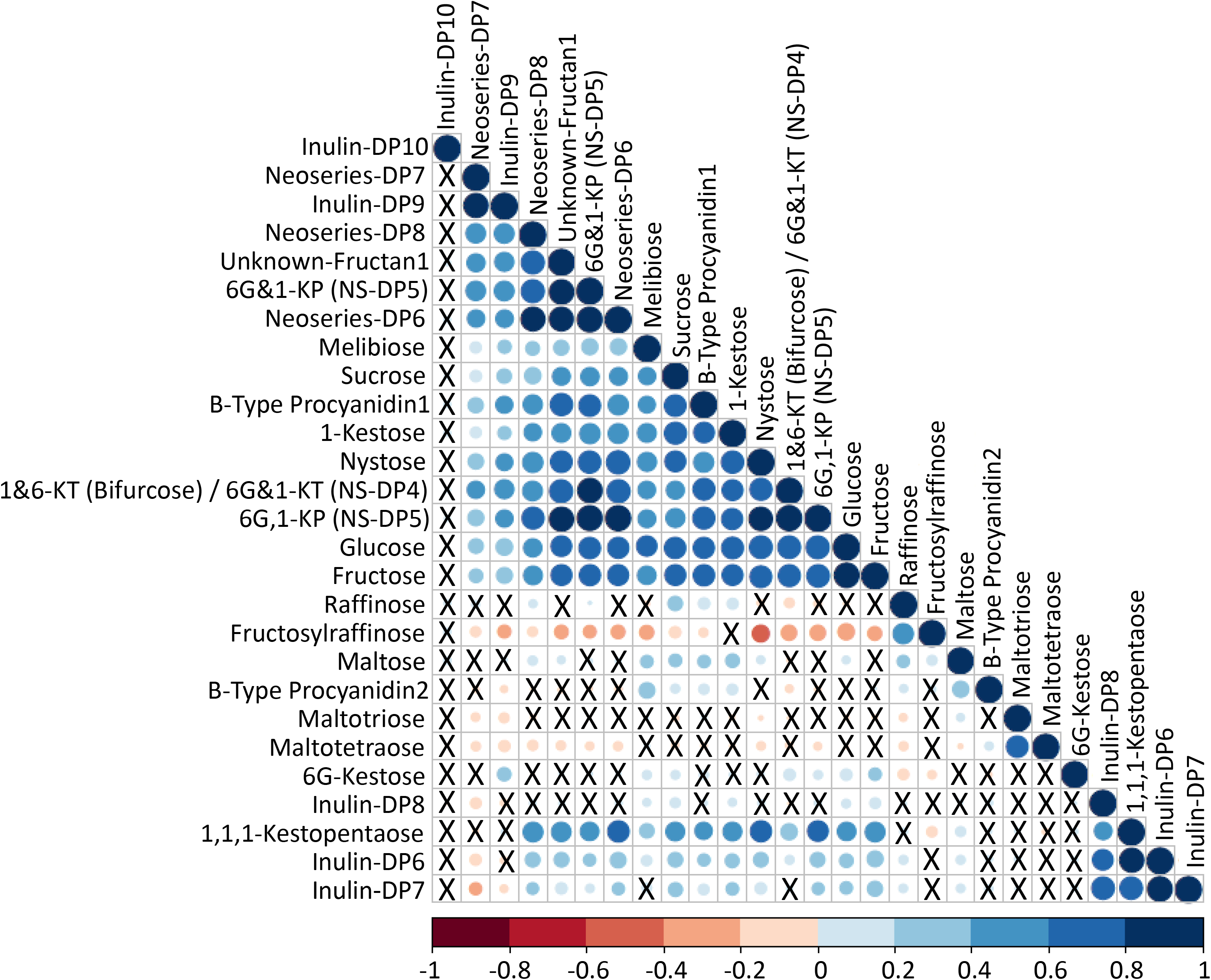
Correlation pattern among metabolites. Pair-wise Pearson correlations are shown in a heat map representation, whereas metabolites are sorted according to correlation-based hierarchical cluster analysis. High positive correlations are represented by dark blue and negative ones by red circles, whereas the circle diameter is indicative for the strength of the correlation. X-not significantly correlated (p>0.05). The raw data are presented in Table S4.

### Differences in grain oligosaccharide profiles are genetically controlled in barley

GWAS for variation in mature grain oligosaccharides identified a single highly significant association for two compounds, neoseries-DP7 (LOD = 8.65, *p* = 2.25 x 10^−9^) and inulin-DP9 (LOD = 6.74, *p* =1.81 x 10^−7^), with other less significant associations for both of these on chromosome 7H (Figure 3A). Regression of these two traits showed a high level of correlation (R^2^ = 0.86, Figure 2, Table S4). Both QTL on 7H overlapped and the most significant marker from the analysis was the same, JHI-Hv50k-2016-438638 (Table 2A). This marker had adjusted p-values after FDR correction of p = 0.00004 for neoseries-DP7 and p = 0.003 for inulin-DP9. We anchored this QTL to the physical map (Mascher *et al.*, 2017), which based on local LD spans 3.88 MB from 174,327 (JHI-Hv50k-2016-435062) to 4,056,691 bases (JHI-Hv50k-2016-439312).

**Figure 3:**
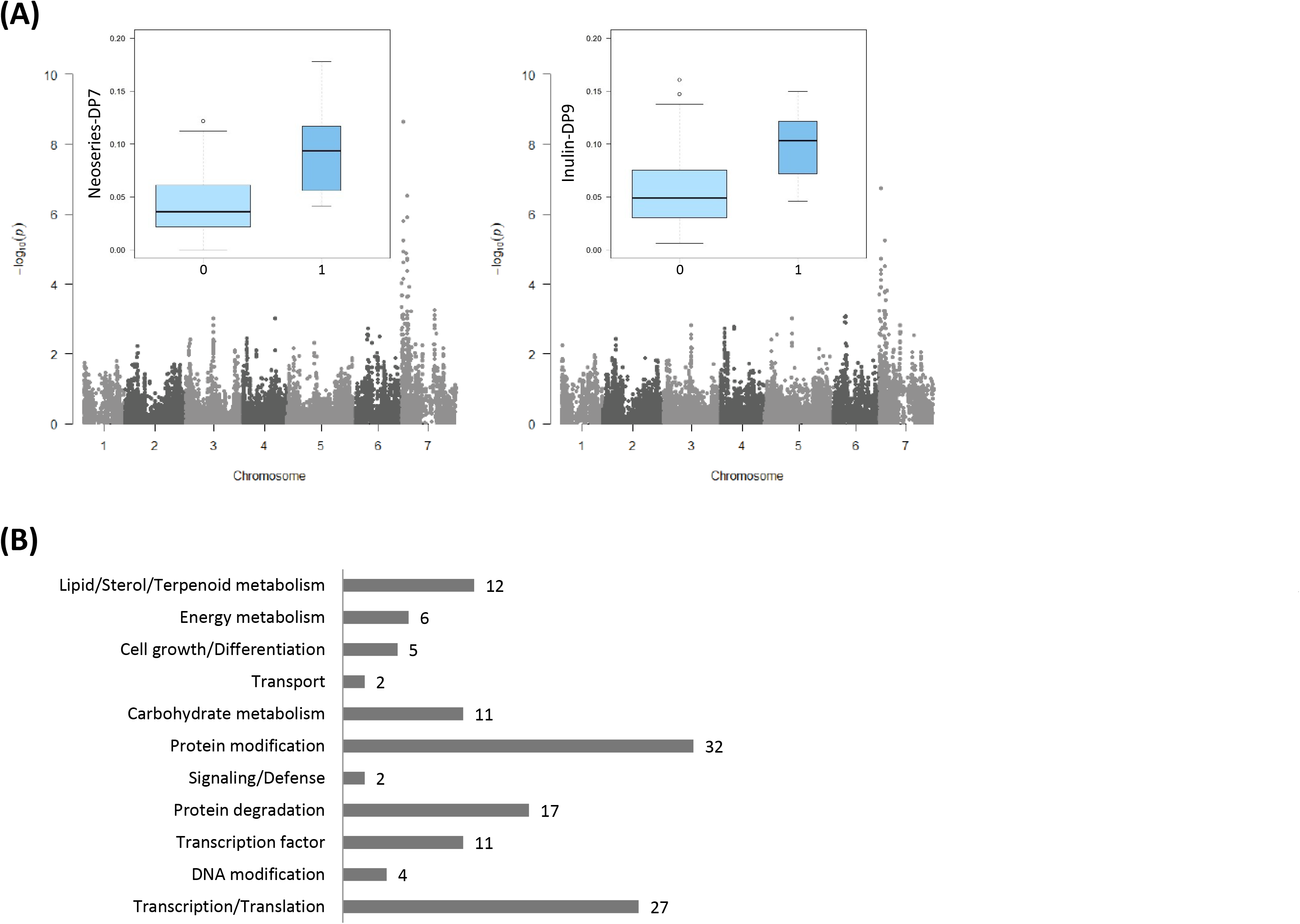
Results of the GWA scan. (A) Manhattan plots are shown for the two fructans neoseries-DP7 and inulin-DP9. The −log10 (p-value) is shown on the y-axis and the 7 barley chromosomes are shown on the x-axis. Marker-trait association analysis was based on mean integrated peak areas. Integrated box plots show information for the top SNP (JHI-Hv50k-2016-438638). The variable alleles found (A and G) are shown on the x-axis. The y-axis shows the median of the metabolite amount for all lines with the respective allele variant; the width of the box is indicative for the number of lines with this particular allele. The false discovery rate significance threshold = −log 10(p) 6.02. (B) Assembling of the annotated gene models from the significant QTL region according to functional categories as obtained from the UniProt database (https://www.uniprot.org/uniprot/, June 2019). Numbers represent the count of gene models with the respective functional annotation.

**Table 2:**
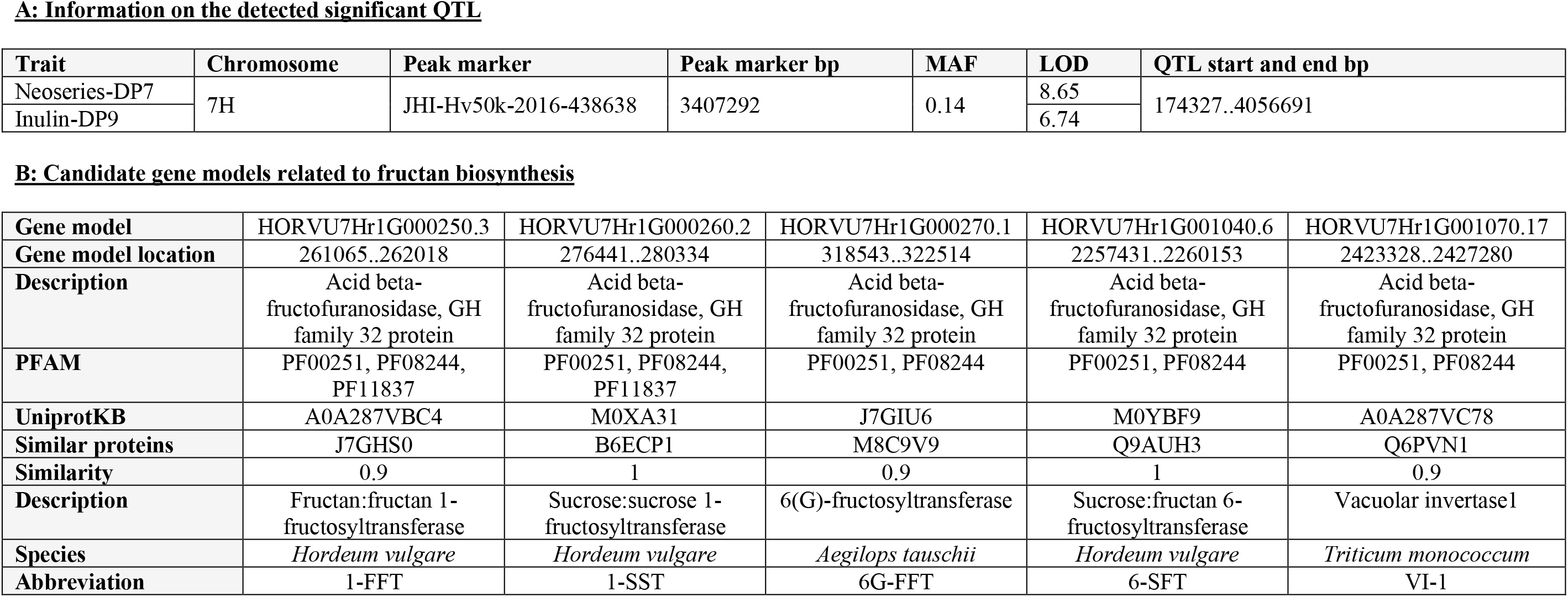
Significant GWA results for the two metabolites neoseries-DP7 and inulin-DP9. Abbreviations: bp, base pair; DP, degree of polymerisation; LOD, logarithm of odds; MAF, minimum allele frequency; QTL, quantitative trait loci

In total 194 gene models were detected within the QTL, of which 65 are unannotated (Table S5). The highest number of annotated gene models was involved in protein modification (32) and degradation (17) (Figure 3B) with others involved in transcription/translation (27), lipid/sterol/terpenoid metabolism (12), transcription factors (TFs) (11), or carbohydrate metabolism (11). Among the latter category, we identified five candidates that could influence fructan content (Table 2B). These included HORVU7Hr1G000250.3, HORVU7Hr1G000260.2, and HORVU7Hr1G001040.6, which were genetically similar or identical to *1-FFT*, *1-SST* and *6-SFT* from *Hordeum vulgare*, respectively. Of the two others, HORVU7Hr1G000270.1 was similar to *6(G)-fructosyltransferase* (*6G-FFT*) from *Aegilops tauschii*, and HORVU7Hr1G001070.17 to *vacuolar invertase1* (*VI-1*) from *Triticum monococcum.*

To further explore relationships between compounds we used a hypothesis-free analysis of metabolite ratios in a GWAS. This analysis generates a ‘p-gain’ statistic which is calculated from the significance of increases in −log10(p) values of the metabolite ratios compared to an estimated threshold derived from the p-values obtained in GWAS of the individual compounds (Petersen *et al.*, 2012). Using the Log transformed ratios between neoseries-DP7 and inulin-DP9 with all other compounds, 17 pairs of compounds correlated with a QTL that passed the FDR threshold of −log 10(p) 6.02 in the same region of chromosome 7H as neoseries-DP7 and inulin-DP9 alone. The ratios of neoseries-DP7:inulin-DP10, inulin-DP9:neoseries-DP8 and inulin-DP9:inulin-DP10 passed the p-gain threshold of 5.2 x 10^2^ (p < 0.05) for markers with a MAF of > 10% with 1.97 x 10^5^, 4.08 x 10^9^ and 3.71 x 10^3^, as well as 1.93 x 10^5^ and 6.43 x 10^5^, respectively (Table 3, Figure S4), indicating metabolic links between these compounds. The QTL on 7H overlapped for all ratios. Significant markers identified were SCRI_RS_8079, JHI-Hv50k-2016-435510, and JHI-Hv50k-2016-438638, the latter being the same as identified with the metabolite concentrations for neoseries-DP7 and inulin-DP9 alone (Table 2A). GWAS for the ratio neoseries-DP7:inulin-DP9 did not identify any significant associations (Table 3).

**Table 3:** Significant GWA results for the ratios of neoseries-DP7 and inulin-DP9 with all other metabolites. Abbreviations: bp, base pair; DP, degree of polymerisation; FDR, false discovery rate; LOD, logarithm of odds; MAF, minimum allele frequency; *P*, probability value; *P*-gain, ratio of the lowest p-value of the two individual metabolites and the p-value of the metabolite ratio; P-values and FDR adjusted p-values from the initial GWAS for traits that did not identify significant associations but when included as a ratio do identify significant associations are included in columns ‘*P* for non sig trait’ and ‘FDR adjusted p-value for non sig trait’; * indicates significant results passing the p-gain threshold of 5.2 x 10^2^, which are also highlighted in light grey. Corresponding Manhattan and box plots are shown in Supplementary Figure S4.

| Ratio combination | Peak marker | Peak marker bp | MAF | LOD | *P* | *P* for non sig trait | FDR adjusted p-value | FDR adjusted p-value for non sig trait | *P*-gain | FDR adjusted p-gain |
| --- | --- | --- | --- | --- | --- | --- | --- | --- | --- | --- |
| Neoseries-DP7:Fructosylraffinose | JHI-Hv50k-2016-438638<br>JHI-Hv50k-2016-438980 | 3407292<br>3689408 | 0.14 | 7.0 | 1.08E-07 | 6.26E-04 | 2.24E-03 | 6.76E-01 | 2.08E-02 | 2.08E-02 |
| Neoseries-DP7:B-Type Procyanidin1 |  |  |  | 6.9 | 1.39E-07 | 4.73E-05 | 2.87E-03 | 1.00E+00 | 1.62E-02 | 1.62E-02 |
| Neoseries-DP7:1,1,1-Kestopentaose |  |  |  | 9.1 | 8.62E-10 | 8.22E-05 | 1.79E-05 | 2.84E-01 | 2.61E+00 | 2.61E+00 |
| Neoseries-DP7:6G,1-KP |  |  |  | 9.0 | 8.96E-10 | 9.06E-01 | 1.86E-05 | 1.00E+00 | 2.51E+00 | 2.51E+00 |
| Neoseries-DP7:Inulin-DP6 |  |  |  | 8.4 | 3.88E-09 | 9.78E-04 | 8.04E-05 | 1.00E+00 | 5.80E-01 | 5.80E-01 |
| Neoseries-DP7:Neoseries-DP6 |  |  |  | 7.9 | 1.28E-08 | 1.51E-01 | 2.66E-04 | 1.00E+00 | 1.75E-01 | 1.75E-01 |
| Neoseries-DP7:Inulin-DP7 |  |  |  | 9.1 | 8.45E-10 | 3.17E-03 | 1.75E-05 | 1.00E+00 | 2.66E+00 | 2.66E+00 |
| Neoseries-DP7:Neoseries-DP8 |  |  |  | 10.8 | 1.64E-11 | 7.89E-02 | 2.14E-07 | 1.00E+00 | 1.37E+02 | 2.17E+02 |
| Neoseries-DP7:Inulin-DP10 | SCRI RS 8079 | 2325562 | 0.11 | 7.4 | 4.24E-08 | 3.46E-01 | 1.72E-03 | 1.00E+00 | 1.97E+05* | 5.80E+02 |
| Inulin-DP9:1-Kestose | JHI-Hv50k-2016-438638<br>JHI-Hv50k-2016-438980 | 3407292<br>3689408 | 0.14 | 8.0 | 1.03E-08 | 1.72E-02 | 2.14E-04 | 1.00E+00 | 1.75E+01 | 1.75E+01 |
| Inulin-DP9:1,1,1-Kestopentaose |  |  |  | 9.4 | 4.38E-10 | 8.22E-05 | 9.08E-06 | 2.84E-01 | 4.14E+02 | 4.14E+02 |
| Inulin-DP9:6G,1-KP |  |  |  | 7.3 | 6.29E-06 | 9.06E-01 | 9.40E-04 | 1.00E+00 | 2.88E-02 | 4.00E+00 |
| Inulin-DP9:Inulin-DP6 |  |  |  | 8.4 | 4.03E-09 | 9.78E-04 | 8.35E-05 | 1.00E+00 | 4.50E+01 | 4.50E+01 |
| Inulin-DP9:Neoseries-DP6 |  |  |  | 8.0 | 1.04E-08 | 1.51E-01 | 2.15E-04 | 1.00E+00 | 1.75E+01 | 1.75E+01 |
| Inulin-DP9:Inulin-DP7 |  |  |  | 9.0 | 9.04E-10 | 3.17E-03 | 1.87E-05 | 1.00E+00 | 2.01E+02 | 2.01E+02 |
| Inulin-DP9:Neoseries-DP8 | JHI-Hv50k-2016-435510<br>JHI-Hv50k-2016-437376 | 280043<br>2350955 | 0.07 | 10.7 | 1.93E-11 | 3.71E-04 | 4.01E-07 | 1.00E+00 | 4.08E+09* | 2.02E+06 |
|  | JHI-Hv50k-2016-438638<br>JHI-Hv50k-2016-438980 | 3407292<br>3689408 | 0.14 | 10.3 | 4.89E-11 | 7.89E-02 | 5.06E-07 | 1.00E+00 | 3.71E+03* | 7.42E+03 |
| Inulin-DP9:Inulin-DP10 | JHI-Hv50k-2016-435510<br>JHI-Hv50k-2016-437376 | 280043<br>2350955 | 0.07 | 6.9 | 1.36E-07 | 3.21E-01 | 1.93E-03 | 1.00E+00 | 1.93E+04* | 4.18E+02 |
|  | SCRI RS 8079 | 2325562 | 0.11 | 6.9 | 1.40E-07 | 3.46E-01 | 1.93E-03 | 1.00E+00 | 6.43E+05* | 5.17E+02 |
| Neoseries-DP7:Inulin-DP9 | JHI-Hv50k-2016-438638<br>JHI-Hv50k-2016-438980 | 3407292,<br>3689408 | 0.14 | 1.6 | 2.31E-02 | 1.80E-07 | 1.00E+00 | 3.74E-03 | 9.75E-08 | 4.66E-05 |

### Evaluation of exome capture data revealed several non-synonymous SNPs

Mascher *et al.* (2017) presented exome capture data for 25 of the genotypes included in our study, which we evaluated to identify putative casual SNPs for our five regional candidates involved in fructan biosynthesis. Eight non-synonymous SNPs in *1-FFT*, three in *VI-1*, two in *6G-FFT*, and one in *1-SST* were identified (Table S6). All identified SNPs are located within functional protein coding regions of the genes (Figure 4). Changes in just one out of 25 genotypes were observed for three markers among the eight SNPs detected in *1-FFT*. The other five SNPs in *1-FFT* represent changes from methionine to leucine (position 7H_ 262685), alanine to threonine (7H_263547 and 7H_263700), isoleucine to threonine (7H_264127), and leucine to isoleucine (7H_264198). They showed significant effects (p < 0.05) on 1-kestose and several neoseries-type fructans (Figure S5). The two SNPs in *6G-FFT* were in LD and represent changes from glycine to glutamic acid (7H_321608), and alanine to threonine (7H_319284). Notably, they have a significant effect on 1-kestose and several inulin-type fructans (Figure S5). However, the SNP in *1-SST*, representing a change from threonine to isoleucine (7H_279526), as well as the three SNPs in *VI-1*, representing changes from glutamic acid to aspartic acid (7H_2423349), tryptophan to arginine (7H_2425560), and arginine to cysteine (7H_2425578), did not show a significant effect on either trait. For *6-SFT* no SNP was identified.

**Figure 4:**
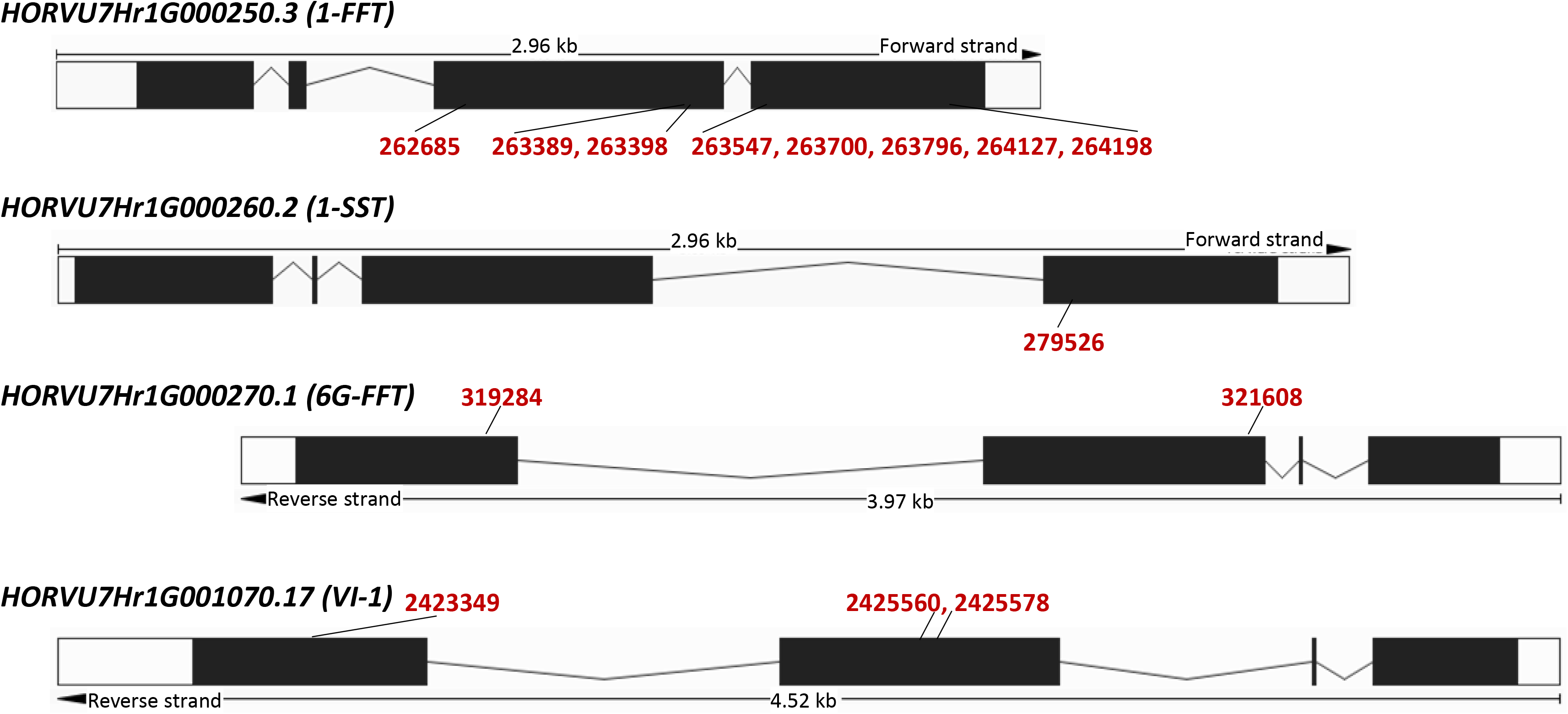
Location of non-synonymous SNPs in fructan biosynthesis candidate genes. All significant causal SNPs (position assigned in bold red) are located within functional protein coding regions of the genes (black regions of the transcripts). All identified SNPs are presented in Table S6 and box plots for significant effects are shown in Figure S5. For *6-SFT* no SNP was identified.

### Fructan biosynthesis genes show developmental stage and tissue specific expression patterns

We compared expression patterns of the five candidate genes (Table 2) and three known fructan hydrolyase encoding genes in various tissues across barley plant development (Figure 5). In the vegetative phase, highest expression for *1-SST* (facilitating the biosynthesis of 1-kestose, the precursor for production of inulin- and graminan-type fructans) was observed during early germination in embryo and all seedling tissues (Figure 5A). During the reproductive phase, *1-SST* is expressed in all vegetative tissues with peak expression in the leaf epidermis, as well as in all reproductive tissues and stages with a pronounced peak of expression in the ovary wall, the embryo sac (ES), the egg apparatus and central cell (EC+CC), and the antipodal cells (ANT) during late pistil development (stages W8 to W10, Figure 5B). In the grain development phase, *1-SST* expression is highest in the early stages (7 to 9 DAP) in maternal grain tissues (pericarp, aleurone, sub-aleurone/outer starchy endosperm (SA)) while decreasing during the storage stage (from 11 DAP onwards) in all tissues (Figure 5C). *1-FFT* (mediating the biosynthesis of inulin-type fructans) showed tight co-expression with *1-SST* during early germination in embryo and all seedling tissues (Figure 5A) as well as all vegetative tissues (Figure 5B), while in reproductive (Figure 5B) and grain tissues (Figure 5C) much lower expression levels were observed. In contrast, *6-SFT* (mediating the biosynthesis of graminans-type fructans) showed very tight co-expression with *1-SST* during meiosis and pistil development (Figure 5B) as well as at early grain development (Figure 5C). During germination, *6-SFT* expression was extremely low while it was observed to be moderate in seedling (Figure 5A) and all vegetative tissues (Figure 5B). Notably, expression of *6G-FFT* (mediating the biosynthesis of neoseries-type fructans) was restricted to the outer grain tissues (see aleurone tissues in Figure 5A and aleurone, pericarp and endosperm tissues in Figure 5C) during late grain development (from 11 DAP onwards). *VI-1*, with yet unknown function, showed low expression levels in germinated grain tissues, all vegetative tissues, in pericarp at late grain development and senescing leaf, while higher levels were notable during late pistil development (Figure 5). Among the fructan hydrolyases, *1-FEH* (HORVU6Hr1g011260) and *6-FEH* (HORVU2Hr1G109120) seem to be involved in balancing fructan biosynthesis, with *1-FEH* tightly co-expressed with *1-SST* in all tissues and stages and pronounced *6-FEH* expression during the reproductive phase in all tissues and in the pericarp at late grain development. In contrast, only marginal expression levels were observed for *6-FEH/CWI2* (HORVU2Hr1G118820) (Figure 5).

**Figure 5:**
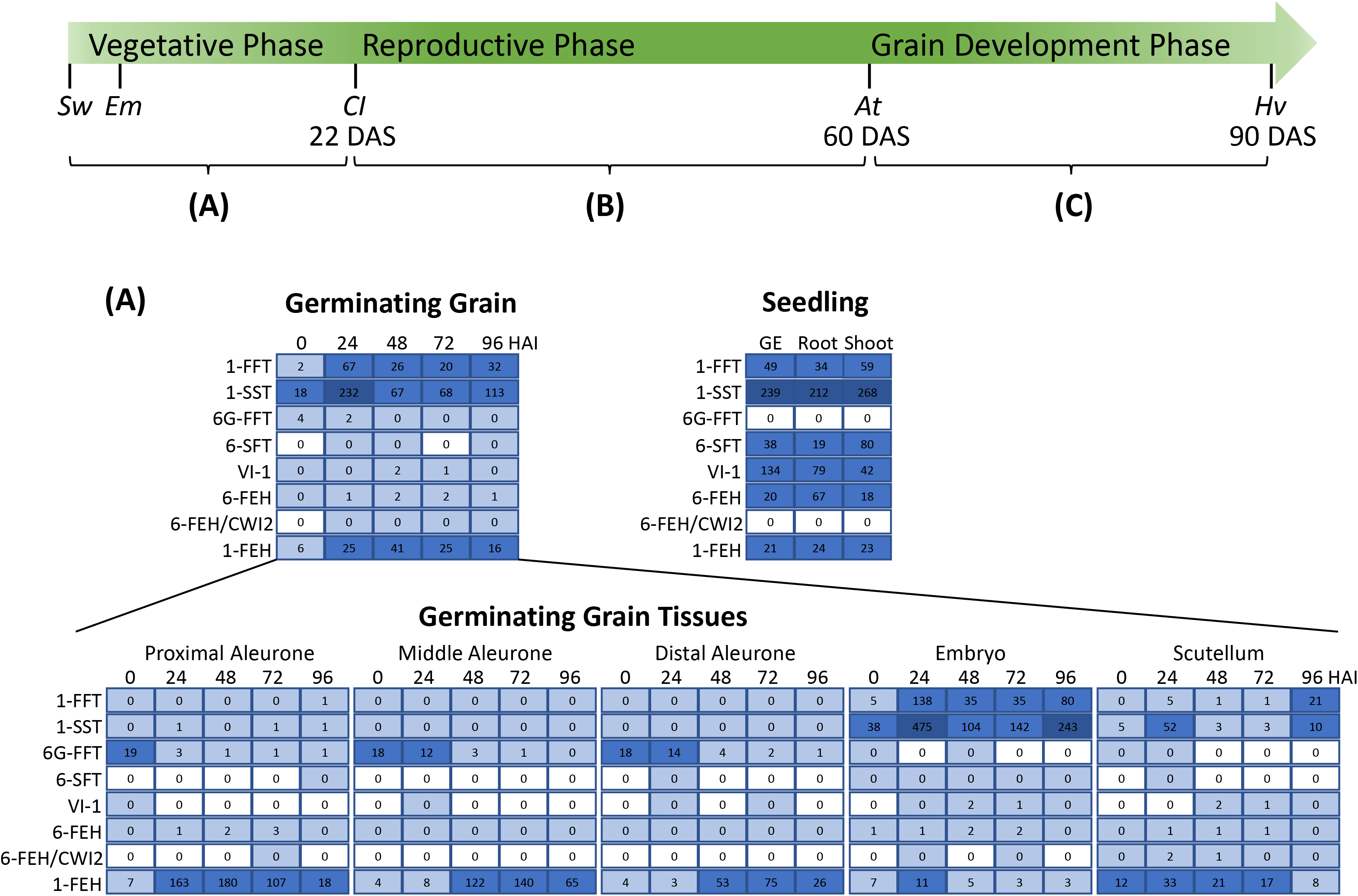
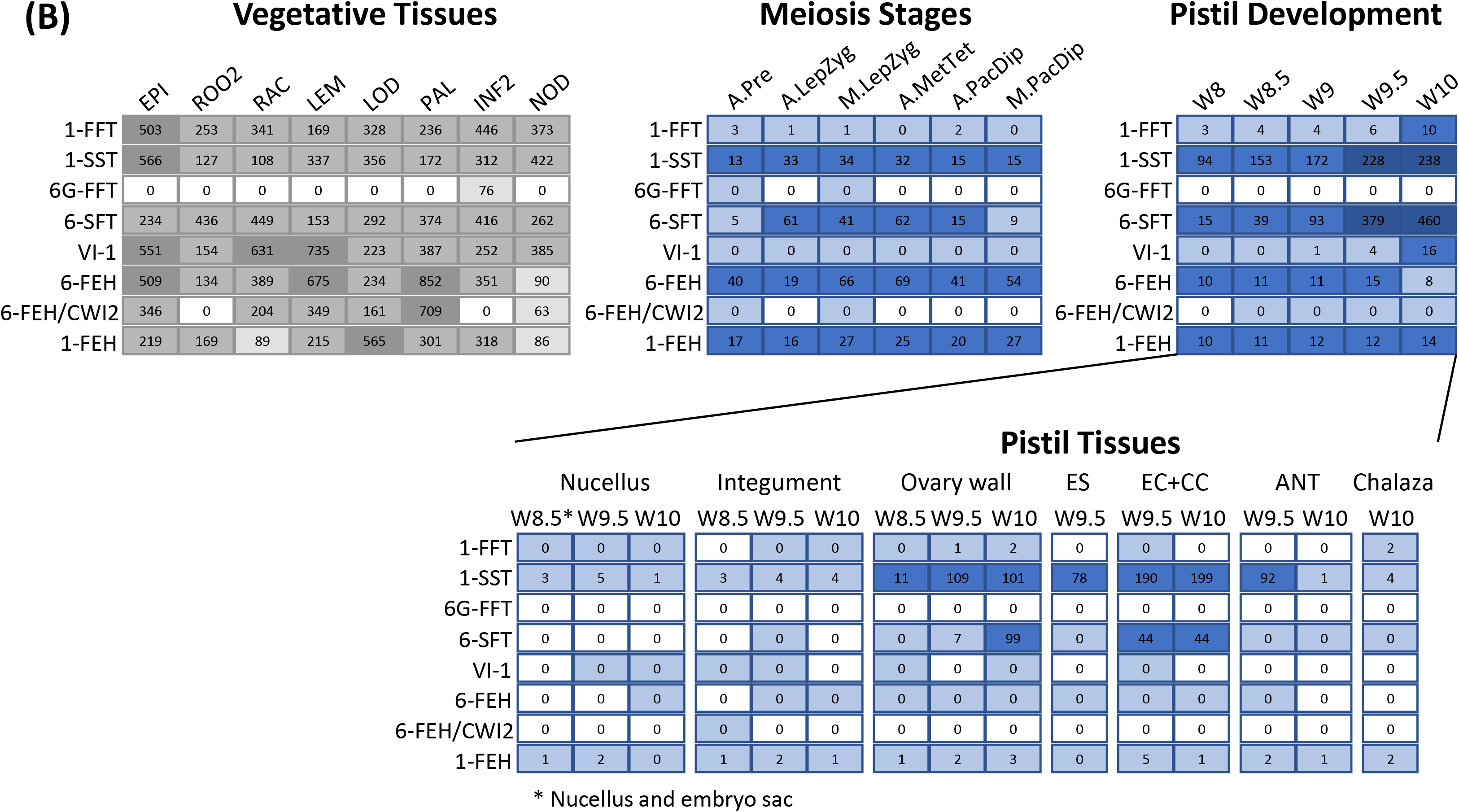
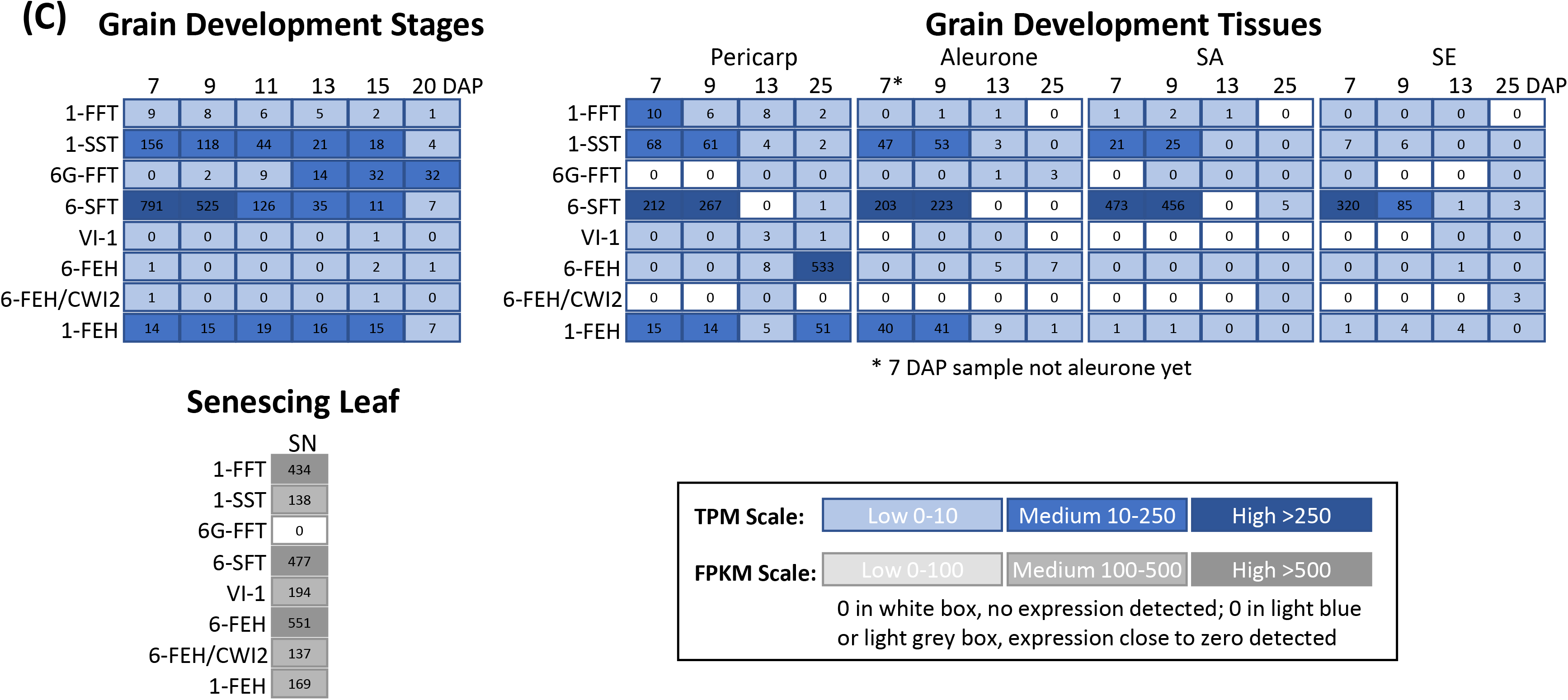
Transcript expression of the fructan biosynthesis genes in barley across plant development and in various tissues. Datasets included in the analysis are detailed in the materials and methods section. Genes included comprise the five fructan biosynthesis genes from the detected significant QTL region (*1-FFT*, HORVU7Hr1G000250; *1-SST*, HORVU7Hr1G000260; *6G-FFT*, HORVU7Hr1G000270; *6-SFT*, HORVU7Hr1G001040; and *VI-1*, HORVU7Hr1G001070) as well as three known fructan hydrolyase encoding genes (*6-FEH*, HORVU2Hr1G109120; *6-FEH/CWI2*, HORVU2Hr1G118820; and *1-FEH*, HORVU6Hr1g011260). Expression levels are colour coded, whereas different scales were used for TPM and FPKM values as indicated in the legend. (A) shows expression levels in the early vegetative phase for whole germinated grain tissues and for isolated germinated grain tissues from 0 to 96 hours after imbibition (HAI). Also, expression levels in seedling are shown for germinated embryo (GE), root and shoot. (B) shows data from the reproductive phase. Vegetative tissues included are: EPI, epidermal strips (4 weeks after planting, W4); ROO2, roots (W4); RAC, inflorescences, rachis (W5); LEM, inflorescences, lemma (W6); LOD, inflorescences, lodicule (W6); PAL, dissected inflorescences, palea (W6); INF2, inflorescence (10 mm); and NOD, internode. Meiosis stages included are: A.Pre, premeiosis anthers; A.LepZyg, leptotene/zygotene anthers; M.Lep/Zyg leptotene/zygotene meiocytes; A.MetTet, metaphaseI-tetrad anthers; A.PacDip, pachytene/diplotene anthers; M.PacDip, pachytene/diplotene meiocytes. Waddington (W) stages for pistil development are: W8; W8.5; W9; W9.5; and W10. Isolated pistil tissues are: nucellus (including nucellus and embryo sac for W8.5); integument; ovary wall; ES, embryo sac; EC+CC, egg apparatus and central cell; ANT, antipodal cells; and chalaza. (C) shows data from the grain development phase for whole developing grain (from 7 to 20 days after pollination, DAP) and for isolated developing grain tissues from 7 to 25 DAP. Abbreviations are SA, sub-aleurone/outer starchy endosperm; and SE, starchy endosperm/inner starchy endosperm. Also, data from senescing leaf (SN) are presented. Other abbreviations are: At, anthesis; CI, collar initiation; Em, emergence; Hv, harvest; and Sw, sowing.

### Fructan biosynthesis genes show differential co-expression patterns in developing barley grain

Besides the five fructan biosynthesis genes, the association of differential oligosaccharide profiles with other candidates in the identified genomic region may be possible. We hypothesised similar expression patterns for fructan biosynthesis genes and other candidates influencing the fructan levels in barley. Therefore, transcript expression levels were evaluated for all gene models within the QTL interval and co-expression of genes was assessed individually within the developmental phases and tissues (Table S7). We have focused on developing barley grain and significant correlations for the expression of fructan metabolism genes with each other and with TFs (Table 4).

**Table 4:**
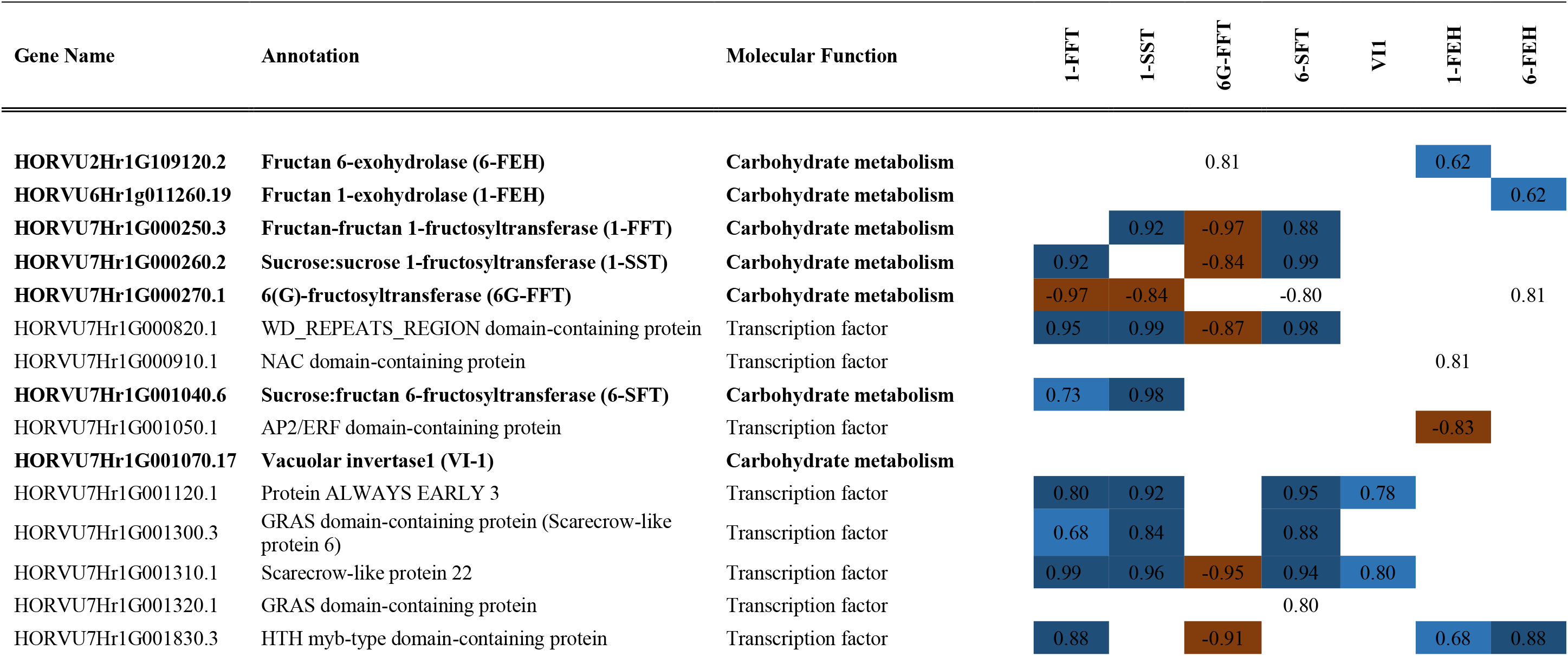
Significant correlations for the expression of fructan metabolism genes with each other and with potential regulatory gene models from the detected QTL interval for developing barley grain. Positive correlations are shown in blue and negative ones in orange, whereas the color code is indicative for the strength of the correlation (the darker, the stronger). Significance threshold was p > 0.05. Numbered boxes without formatting indicate values just above significance (p < 0.055). Bold framed boxes indicate correlations which were detected in various developing grain datasets. Datasets correspond to the ones presented in Figure 5 and methods are detailed in section ‘material and methods’. The raw data are presented in Table S8.

Highly positive correlations among the fructan metabolism genes were observed between *1-FFT*, *1-SST* and *6-SFT*; between *6G-FFT* and *6-FEH*; and for *1-FEH* with *1-FFT*, and *6-FEH*. Notable negative correlations were observed for *6G-FFT* with *1-FFT*, and *1-SST*. Co-expression patterns with TFs were highly similar for *1-FFT*, *1-SST* and *6-SFT* in the developing grain. Notable positive correlations for *1-FFT*, *1-SST* and *6-SFT* expression were identified with the *WD_REPEATS_REGION domain-containing protein* (HORVU7Hr1G000820.1), the *ALWAYS EARLY 3* (HORVU7Hr1G001120.1), and the two *scarecrow-like protein* genes (HORVU7Hr1G001300.3, HORVU7Hr1G001310.1). In contrast, *6G-FFT* showed significant negative correlations with the *WD_REPEATS_REGION domain-containing protein* (HORVU7Hr1G000820.1), *scarecrow-like protein 22* (HORVU7Hr1G001310.1), and *HTH myb-type domain-containing protein* (HORVU7Hr1G001830.3). Significant positive correlations for *VI-1* were observed in developing grain with the *protein ALWAYS EARLY 3* (HORVU7Hr1G001120.1) and *scarecrow-like protein 22* (HORVU7Hr1G001310.1), as observed for *1-FFT*, *1-SST*, and *6-SFT*. *1-FEH* showed co-expression patterns partly like those observed for *1-FFT*. Besides the positive correlation with a *myb-type transcription factor* (HORVU7Hr1G001830.3) a strong negative correlation with the *AP2/ERF domain-containing protein* (HORVU7Hr1G001050.1) and a positive interaction (not significant) with a *NAC domain-containing protein gene* (HORVU7Hr1G000910.1) were identified. Highest positive correlations were noted for *6-FEH* with *6G-FFT* and the *HTH myb-type domain-containing protein* (HORVU7Hr1G001830.3), which in contrast was negatively associated with *6G-FFT* (Table 4).

Additional and partly different patterns for the co-expression of fructan metabolism genes with other genes were observed across a range of developmental phases and tissues (Table S7, Supplementary results).

## Discussion

### Neoseries-type fructans are abundant in mature barley grain

Profiling of DP3 to DP10 oligosaccharides revealed the abundance of 6G-kestose and higher DP neoseries-type fructans in mature barley grain (Table 1, Figure S1, Table S2). While the presence of 6G-kestose has been reported in wheat and barley grain (Nilsson *et al.*, 1986; Henry and Saini, 1989) higher DP variants of this fructan-type have not been previously identified. Recent studies revealed the presence of neoseries-type fructans in oat, rye, spelt and wheat flour (Verspreet *et al.*, 2015b; Verspreet *et al.*, 2017) but claimed its absence in barley (Verspreet *et al.*, 2017). These contrasting observations may be explained both by the plant materials used (flour vs. whole grain) and the technical constraints of fructan profiling. Noticeable accumulation of unidentified higher DP fructans was reported for the outer pericarp of developing wheat grain (Schnyder *et al.*, 1993). Thus, utilising whole grain here may have facilitated the detection of neoseries-type fructans in barley, likely accumulating in outer grain parts as suggested by expression analysis for *6G-FFT* (Figure 5). Additionally, electronic properties of PAD, typically used for fructan profiling, require higher concentrations for the detection of higher molecular weight fructans (Rocklin and Pohl, 1983), resulting in a pronounced log-logistic distribution for those compounds that is also observed in our study (Figure S2) and which may have led to the underrepresentation of fructans with >DP4 in other studies. In the future, comprehensive grain fructan profiling could be improved by employing recently established LC-MS methodologies, as reviewed in Matros *et al.* (2019).

### Barley accessions group according to their oligosaccharide accumulation patterns

Reported genotypic variation in grain fructan content ranges from 0.9-4.2% of dry matter (DM) among 20 barley breeding lines (Nemeth et al., 2014) and from 1.1-1.6% of DM among seven barley cultivars (De Arcangelis *et al.,* 2019). These results correspond well with the variability for total fructan values (0.02-1.94% of grain dry weight) determined here. When discriminating between different chain lengths Henry and Saini (1989) measured varying amounts of FOS with 0.26% (DP3), 0.2% (DP4), 0.03% (DP5) and 0.23% of DM (>DP5) in mature barley grains, which was confirmed by results from Jenkins *et al.*, (2011). In our study, the lowest abundance range showed FOS with DP5 (traces to 0.16% of DM) while FOS with DP3 and DP4 ranged from 0.02-0.53% and traces to 0.44% of DM, respectively. Nemeth *et al.*, (2014) observed a positive correlation between fructan values and the content of long chain fructans (> DP9, r = 0.54, p = 0.021). However, such an association could not be found in our dataset. Clustering of the oligosaccharide profiles from the 154 lines revealed two major profile groups, one each of higher and lower sugar values (Figure 1). We detected significant positive correlations between biosynthetically closely related metabolites (e.g. within and between the different fructan-types) with negative associations for antagonistic compounds (e.g. fructosylraffinose with all fructans, sugar monomers and dimers, Figure 2). Co-occurrence of fructans and RFO has been reported for many plant species including wheat (Haska *et al.*, 2008) and barley (Henry, 1988), with the proposal that strong RFO and fructan accumulation do not occur together in a single plant species (Van den Ende, 2013). Notably, fructosylraffinose was only speculated to occur in barley (Cerning and Guilbot, 1973), while its presence was described in wheat decades ago (White and Secor, 1953; Saunders, 1971).

### Differences in oligosaccharide profiles are genetically controlled in barley

A significant QTL on chromosome 7H affecting barley grain fructan levels was identified (Figure 3A) and five genes involved in fructan metabolism were detected in this region (Table 2 and Table 3, Table S5). We increased the power of GWAS by analysing metabolite ratios using the p-gain approach. As the p-gain passed an appropriate threshold (defined by the data), using ratios provided more information about the traits, and the genomic locus underlying them, than looking at the traits individually. Using ratios reduces background ‘noise’ in datasets, increasing statistical power to detect significant associations between traits and genomic loci (Petersen *et al.*, 2012). Previous studies have demonstrated that including ratios between pairs of traits can strengthen associations identified and uncover novel information about biochemical pathways (Gieger *et al.*, 2008; Illig *et al.*, 2010; Suhre *et al.*, 2011). Thus, ratio-GWAS represents an innovative approach for the discovery of new biologically meaningful associations in plants, as shown for the linked oligosaccharide pathways described here. However, when we used the ratio between neoseries-DP7:inulin-DP9 in the GWAS we did not identify an association on 7H, indicating less information provided by the ratios than the individual values. This may relate to the high positive correlation of these two compounds across the barley lines (R^2^ = 0.86, Figure 2, Table S4) and the close genomic location of the related fructan biosynthesis genes (Table 2). In contrast, for the ratios between inulin-DP7:inulin-DP10 (R^2^ = 0.011, p = 0.80), inulin-DP9:neoseries-DP8 (R^2^ = 0.56, p< 0.05) and inulin-DP9:inulin-DP10 (R^2^ = 0.028, p = 0.54) a significant QTL was identified. This points towards a stronger association between the identified genomic locus and the molecular weight of the fructans than with the fructan structure.

In wheat, two loci for differential total fructan contents in grain were identified on chromosomes 7A and 6D, which did not show significant interactions (Huynh *et al.*, 2008b). Subsequent physical mapping provided indications for clustering of fructan biosynthesis genes in the genomes of both dicots as well as monocots (Huynh *et al.*, 2012). For wheat and barley the formation of a functional cluster was shown containing *1-SST* (provided are the IDs of the most probable barley gene product; J7GM45_HORVV), *1-FFT* (J7GHS0_HORVV), and *6-SFT* (Q96466_HORVU) (Huynh *et al.*, 2012), which were also identified here. Additionally, the authors found two *vacuolar invertase* (*VI*) genes (J7GIU6_HORVV, J7GR98_HORVV) in this cluster, of which we identified one, which is similar to *6G-FFT* (J7GIU6_HORVV). The identification of *6G-FFT* matches the detection of neoseries-type fructans in our study. Among the five candidates we identified was also a gene coding for an uncharacterised gene product (M0X3V0_HORVV) which is similar to a *VI-1* from *T. monococcum* (Q6PVN1_TRIMO) that has not been described or annotated in barley before.

The evaluation of exome capture data (Mascher *et al.*, 2017) led to the identification of several significant SNPs in the five fructan biosynthesis genes. SNPs in *1-FFT* were associated with grain neoseries-type fructan content while SNPS in *6G-FFT* were associated with inulin-type fructan content (Figure 4, Table S6). Ideally, the influence of these SNPs would be validated in the complete set of germplasm used to quantify fructan content. This analysis would likely reveal additional SNPs that have not been identified in this subset of lines.

### Developmental and tissue specific nature of barley fructan biosynthesis

Throughout plant development, *1-SST*, *1-FFT* and *1-FEH* showed strong co-expression, starting in embryo tissue during germination, accompanied later by *6-SFT* expression in root, leaf and stem, likely leading to the biosynthesis of inulin- and graminan-type fructans in those tissues until senescence (Figure 5). These observations matched the consensus of inulin- and graminan-type fructans being the predominant polymers in barley tissues (Pollock and Cairns, 1991; Bonnett *et al.*, 1997; Huynh *et al.*, 2008a). Accordingly, *1-SST*, *1-FFT*, *6-SFT* and *1-FEH* are the best studied fructan biosynthesis genes (Duchateau *et al.*, 1995; Henson, 2000; Lüscher *et al.*, 2000; Huynh *et al.*, 2012). A key role was assigned to *1-SST* (Wagner *et al.*, 1983) and correlated transcription and activity was reported for *1-SST* and *6-SFT* in barley leaves (Nagaraj *et al.*, 2004). A role for fructans as a temporal carbohydrate reserve has been widely accepted in vegetative tissues and roots (Pollock *et al.*, 1996; Vijn and Smeekens, 1999; Housley, 2000) and can be assumed for the inulin and graminan-type fructans in barley.

We observed co-expression of *1-SST*, *6-SFT*, *6-FEH*, and *1-FEH* in reproductive tissues with a pronounced peak during late pistil development in ovary tissues (Figure 5B) probably leading to specific accumulation of graminan-type fructans. In *Campanula rapunculoides*, the largest inulin-type fructan concentrations were found in petals and ovaries (Vergauwen *et al.*, 2000). Based on the observation that petals in daylily (*Hemerocallis*) (Bieleski, 1993) and *C. rapunculoides* (Vergauwen *et al.*, 2000) and leaves of *Phippsia algida* (Solhaug and Aares, 1994) rapidly degrade fructans upon flower opening, a role for them in flower expansion was suggested. However, the function of the different fructan-types accumulating in *C. rapunculoides* ovary (inulin-type) and barley ovary tissues (graminan-type) remains unresolved at present. Also, the newly identified *VI-1* showed peak expression specific to ovary tissues at late pistil development, while its function in fructan biosynthesis remains unclear.

In accordance with the detection of neoseries-type fructans and the identification of *6G-FFT* in the significant QTL region we observed *6G-FFT* expression in barley grain (Figure 5A and C). Notably, its expression was restricted to developing grain from 11 DAP onwards and confined to the outer endosperm and maternal tissues. Reports on this fructan type in developing grains of other cereals do not exist to our knowledge. Accumulation of neoseries-type fructans in the aleurone of mature grain may be related to favourable structural characteristics when compared to inulins and graminans and to the function of this tissue during germination. Some reports showed that fructan branching architecture is critical to physicochemical properties, such as water solubility or formation of aggregates at high concentration (Eigner *et al.*, 1988; Wolff *et al.*, 2000; Ponce *et al.*, 2008). The more compact shape of neoseries-type fructans would allow higher concentrations to be stored in the desiccated aleurone. Better water solubility and pH-stability of neoseries-type fructans would be advantageous during germination, when the aleurone hydrates and enzymes must be quickly activated and reach their substrates. Proving these hypotheses will require the comparative evaluation of physicochemical properties of neoseries-inulin- and graminan-type fructans in the future. Indeed, several studies have provided strong evidence for a positive relationship between enhanced fructan concentrations with better malting characteristics in barley varieties (Smith *et al.*, 1980; Cozzolino *et al.*, 2016 and references therein)

### Potential regulators of barley grain fructan biosynthesis

Despite the increasing evidence of tissue specificity, there is limited knowledge of how fructan metabolism is orchestrated to adjust the storage and use of photosynthates during grain development. Within the significant QTL interval, we found several genes differentially co-expressed with the various fructan biosynthesis genes in developing grain. Among them were several TFs (Table 4, Table S7, Figure 6).

**Figure 6:**
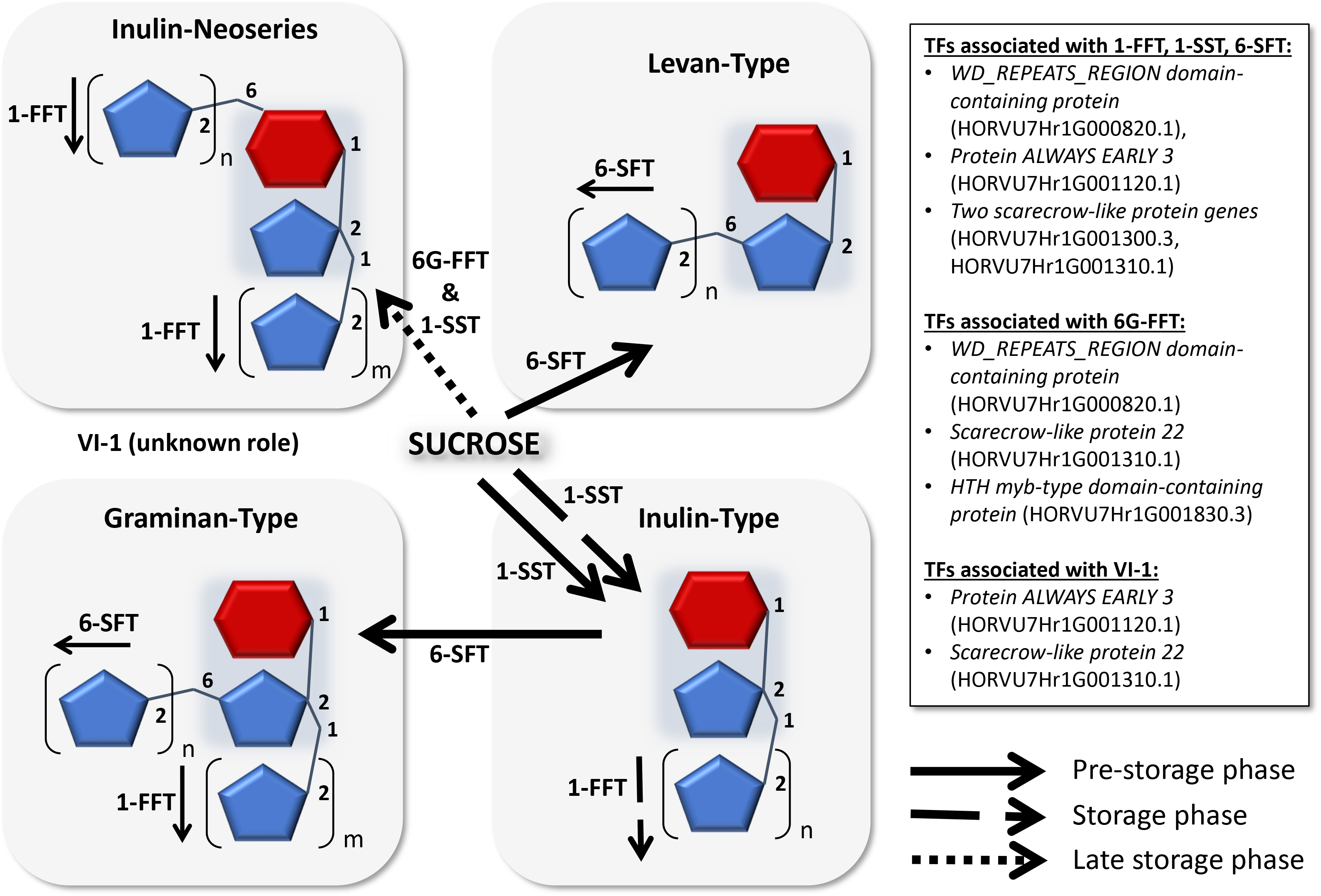
Fructan-types, suggested biosynthesis routes and potential regulators in developing barley grain. Specific spatiotemporal biosynthesis of oligofructans was observed for barley grains. Continuous bold arrows illustrate the major route of biosynthesis during the pre-storage phase (until 14 DAP) and the dashed bold arrows indicate the major route during the storage phase (until 20 DAP). During the pre-storage phase high transcript levels for *1-SST* and *6-SFT* were observed for the endosperm leading to an accumulation of 6-kestose and bifurcose. With transition to the storage phase a transcriptional switch was observed resulting in high transcript levels of *1-SST* in the nucellar projection (NP). *1-FFT* was found to be exclusively expressed in the NP during the storage phase. Induction of the inulin-type fructan biosynthesis pathway led to high amounts of 1-kestose and nystose accumulating in the endosperm cavity (Peukert et al., 2014). The dotted bold arrow illustrates the major route of biosynthesis during the late storage phase with *6G-FFT* transcripts detected in the outer endosperm (from 30 DAP onwards, Figure 5), which matched the detection of neoseries-type oligofructans in mature barley grains (Figure 1). Transcription factors (TF) showing significant correlation of transcript expression pattern in developing grain with *1-FFT, 1-SST* and *6-SFT* (positive), *6G-FFT* (negative), and *VI-1* (positive) are listed in the inserted text box. Inulin-neoseries represents linear fructans with β(2,1) & β(2,6) linked fructosyl units at the glucose (1F, 6G-di-β-D-fructofuranosylsucrose is shown; m=1, n= 1), graminan-type represents branched fructans with β(2,1) & β(2,6) linked fructosyl units (bifurcose is shown; m= 1, n= 1), inulin-type illustrates linear fructans with β(2,1) linked fructosyl units (1-kestose is shown; n= 1);, and levan-type shows linear fructans with β(2,6) linked fructosyl units (6-kestose is shown; n= 1). The arrows indicate direction of further polymerisation. Abbreviations are: *1-FFT*, fructan:fructan 1-fructosyltransferase; *1-SST*, sucrose:sucrose 1-fructosyl-transferase; *6-SFT*, sucrose:fructan 6-fructosyltransferase; *6G-FFT*, fructan:fructan 6G-fructosyltransferase; *VI-1*, vacuolar invertase 1 (unknown role in developing grain).

It is generally agreed that initiation of fructan biosynthesis is triggered by an organ-specific sucrose threshold (Lu *et al.*, 2002, Jin *et al.*, 2017). Also, several molecular components in sucrose-mediated induction of plant fructan biosynthesis, such as protein phosphatases and kinases (Noël *et al.*, 2001), second messenger Ca^2+^ (Martinez-Noël *et al.*, 2006), small GTPases and phosphatidylinositol 3-kinase (Ritsema *et al.*, 2009), as well as the plant hormones abscisic acid and auxin (Valluru, 2015) were shown to be required for activation of fructosyltransferase genes.

An opposing sugar-sensing system was recently identified in barley, whereby a single gene on chromosome 2H encodes two functionally distinct TF variants [SUSIBA (sugar signaling in barley) 1 and 2], which respond differently to sucrose concentrations (Jin *et al.*, 2017). However, no distinction was made between different tissues and fructan types and it remains unclear if this system coordinates fructan and starch biosynthesis in general.

In wheat, TaMYB13, a R2R3-MYB TF, was described as a transcriptional activator of fructan biosynthesis (Xue *et al.*, 2011; Kooiker *et al.*, 2013). In the promoter of the barley genes *1-FFT*, *1-SST*, *6-SFT* and *VI,* binding motifs for TaMYB13 were identified, suggesting that the co-expression of these genes may be driven by a TaMYB13 homolog (Huynh et al., 2012). However, the *HTH myb-type domain-containing protein* (HORVU7Hr1G001830.3) identified here did not show similarity to TaMYB13 and we could not observe a clear homolog in barley. While three myb-type TFs were also described to activate promoters of genes involved in fructan biosynthesis and degradation in chicory (Wei *et al.*, 2017a and b), involvement of other TF family genes has not yet been reported.

## Conclusions

A new genomic region and several causal SNPs involved in the regulation of barley grain fructan content were identified. The genomic region includes a physical cluster of functionally related fructan biosynthetic genes and several potential regulatory genes. While the clustering of fructan biosynthetic genes may hint at the co-evolution of these gene families, a conserved gene co-expression suggesting an equal contribution to grain fructan biosynthesis was not observed. Instead the spatiotemporal dynamics for fructan biosynthetic genes point towards versatile roles of the different fructan types. Phylogenetic relationships between fructosyltransferases and invertases within *Poaceae* suggest that *6-SFT* may have evolved from a *Poaceae* ancestor genome after the major clade of vacuolar invertases diverged, followed then by *1-FFT* and *1-SST* (Huynh *et al.*, 2012). The analysis also showed the presence of a unique barley clade of four vacuolar invertase genes, among them the newly annotated *6G-FFT*, between the *6-SFT* and the *1-FFT* and *1-SST* clades, suggesting that extra duplication might have occurred in barley. Accordingly, in developing grain we observed similar co-expression with a set of TFs for *1-FFT*, *1-SST*, and *6-SFT*, which was different from the associations found for *6G-FFT*. The proposed dynamics of fructan biosynthesis in barley grain and potential regulators are presented in Figure 6. Assuming a specific spatiotemporal control of grain fructan biosynthesis, breeding or genetic engineering for high fructan content related to grain specific traits (e.g. nutritional quality or germination) will require careful approaches targeting certain tissues and developmental stages as recently suggested for engineering mixed linkage (1,3;1,4)-β-glucan biosynthesis in the endosperm (Lim *et al.*, 2019).

## Abbreviations

1-FFT: fructan:fructan 1-fructosyltransferase
1-SST: sucrose:sucrose 1-fructosyltransferase
6-SFT: sucrose:fructan 6-fructosyltransferase
6G-FFT: 6(G)-fructosyltransferase
DAP: days after pollination
DP: degree of polymerisation
DM: dry matter
ELSD: evaporative light scattering detection
FDR: false discovery rate
FOS: fructooligosaccharides
FPKM: fragments per kilobase, per million mapped reads
GWA: genome wide association
GWAS: genome wide association study
HAI: hours after imbibition
HPAEC–PAD: high pH anion exchange chromatography with pulsed amperometric detection
HPLC: high performance liquid chromatography
KP: kestopentaose
KT: kestotetraose
LC: liquid chromatography
LD: linkage disequilibrium
LOD: logarithm of odds
MAF: minimum allele frequency
MS: mass spectrometry
NG: Neural Gas
NS: neoseries-type fructan
P: probability value
PEG: polyethylene glycol
QTL: quantitative trait loci
RFO: raffinose family oligosaccharides
RT: retention time
SNP: single nucleotide polymorphisms
SPE: solid phase extraction
TFA: trifluoroacetic acid
TPM: transcripts per million
VI-1: vacuolar invertase1

## Supplementary data

**Supplementary Table S1:** List of germplasms with their corresponding oligosaccharide profile group.

**Supplementary Table S2:** Oligosaccharide annotation information.

**Supplementary Table S3:** Oligosaccharide peak area entry means values.

**Supplementary Figure S1:** Representative chromatogram of mature barley grain soluble carbohydrates

**Supplementary Figure S2:** Metabolite distribution among the lines.

**Supplementary Figure S3:** Oligosaccharide profile prototypes as obtained by Neural Gas clustering.

**Supplementary Table S4:** Correlation and significance values for associations among the 27 metabolites.

**Supplementary Figure S4:** Manhattan plots and box plots for all significant results from the ratio GWAS.

**Supplementary Table S5:** List of all detected gene models in the significant QTL interval.

**Supplementary Table S6:** Results from exome capture data evaluation.

**Supplementary Figure S5:** Box plots for significant effects of SNPs identified from exome capture data.

**Supplementary Table S7:** Heatmap of correlations for the expression of fructan genes with other gene models from the QTL interval for all developmental phases and tissues.

**Supplementary Table S8:** Expression data and Pearson correlation values for associations among the gene models from the GWA scan for all developmental phases and tissues.

## Acknowledgements

This work was supported by grants from the German Research Foundation (DFG, MA 4814/3-1 & 3-2) to Andrea Matros and the Australian Research Council (ARC) Centre of Excellence in Plant Energy Biology (http://www.plantenergy.uwa.edu.au/) to Rachel Burton. Kelly Houston and Robbie Waugh acknowledge the support of the Rural & Environment Science & Analytical Services Division of the Scottish Government We greatly acknowledge the excellent technical assistance of Shi F. Khor, Jelle Lahnstein. We wish to thank Natalie S. Betts and Helen M. Collins for their help with the geminated grain dataset. The authors acknowledge the use of the facilities, and scientific and technical assistance of the Australian Plant Phenomics Facility, which is supported by the Australian Government’s National Collaborative Research Infrastructure Strategy (NCRIS).

## Author Contributions

AM and RAB designed and developed the concept of the study. RW provided the material and genomic data of the barley panel and was involved in the design of the GWAS. AM and BB conducted the growth experiments and harvested the mature grain material at TPA. AM performed the oligosaccharide profiling analysis and evaluated the HPAEC-PAD data. AM and KW conducted the experiments related to the identification of fructan structures (isolation and MS identification). US performed the clustering of the data and the distribution analysis. KH performed the GWAS and ratio-GWAS as well as the evaluation of the exome capture data. MRT, MKA and LGW conducted the transcriptomic analyses of pistil tissues and developing grain tissues. MS and RW conducted the transcriptomic analysis of developing anther tissues. AM evaluated the transcript expression datasets provided and conducted the co-expression analyses. AM, KH and RAB wrote and provided a draft manuscript which has been revised and accepted by all authors.

